# HandyCNV: Standardized Summary, Annotation, Comparison, and Visualization of CNV, CNVR and ROH

**DOI:** 10.1101/2021.04.05.438403

**Authors:** Jinghang Zhou, Liyuan Liu, Thomas J. Lopdell, Dorian J. Garrick, Yuangang Shi

## Abstract

Here we present an R package for summarizing, annotating, converting, comparing and visualizing CNV (copy number variants) and ROH (runs of homozygosity) detected from SNP (single nucleotide polymorphism) genotyping data. This one-stop post-analysis system is standardized, comprehensive, reproducible, timesaving and user friendly for research in humans and most diploid livestock species.

**Availability and Implementation:** The source code, demo data and vignettes can be found at https://github.com/JH-Zhou/HandyCNV

## 1. Introduction

As the cost of SNP genotyping continues to decrease, a large number of individuals with genome-wide data have been accumulated in the study of various species. In addition to using these data to do GWAS (genome wide association study) or GS (genomic selection), there is more interesting genomic information we can explore from the CNV and ROH that can be inferred from the genotypes. A range of software products have been developed to detect CNV and ROH for SNP data, but few tools have focused on the integration of the detailed summaries, annotations, comparisons, and visualizations of these results. Exploring the useful information from CNV and ROH is time consuming, especially when processing multiple results from different models and software. In order to get more comprehensive results, researchers might often switch back and forth between different tools, and in that case it is easy to make mistakes.

Here we summarized some common unclear or incorrect statements in the literature related to CNV analysis. The most ambiguous statement is when reporting candidate genes in a CNVR (copy number variation region) without the frequency of copy number variation in the genes. Some lower frequency genes (such as only one or two samples exhibiting copy number variation in those genes) might get undue attention in that situation. The second issue is when comparing CNVs between different results and comparisons are made only at population level and ignoring the individual level. Comparison at the population level could reflect the repeatability of CNVs, but at the individual level will examine the similarity between the results. The third issue is when comparing the CNVRs with different reference genomes by converting the coordinates without quality control on the new converted intervals. Quality control is an important step, because the order of SNPs on chromosomes might differ among different reference assemblies, so that the conversion of position for intervals will change the total length of the newly converted intervals, which might cause meaningless comparison. The fourth issue is also common in comparing CNVRs, some literature presented the overlapped number of CNVR via Venn diagram without special illustration. The overlapped number is relative to the results and can spread some confusing information because some longer intervals might overlap multiple shorter intervals from another result, in this case, the overlapped numbers are different relative to each result.

There are several common demands while processing CNV and ROH results, such as preparing summary tables, making summary figures, generating CNVR, plotting CNVR distribution maps, annotating genes, comparing CNVs, comparing CNVRs, converting coordinates, converting map files, finding high frequency abnormal genomic regions, getting consensus genes and custom visualization of results, getting haplotypes of samples in an interested region, *et al*.

There are some small details that are easily overlooked and there are many common analysis steps in post-analysis of CNV and ROH. Therefore, we built this open source tool to provide a standardized, reproducible, time-saving and widely available one-stop post-analysis system to make research more simple, practical and efficient.

## 2. Main Feature

The most useful features provided are: integrating summarized results, generating lists of CNVR, annotating the results with known gene positions, plotting CNVR distribution maps, and producing customised visualisations of CNVs and ROHs with gene and other related information on one plot (Fig. 1). This package supports a range of customisations, including the colour, size of high resolution figures, and choice of output folder to avoid conflict between the results of different runs. Some output files are compatible with other software such as PennCNV, Plink or David annotation tools.

**Fig. 1.**
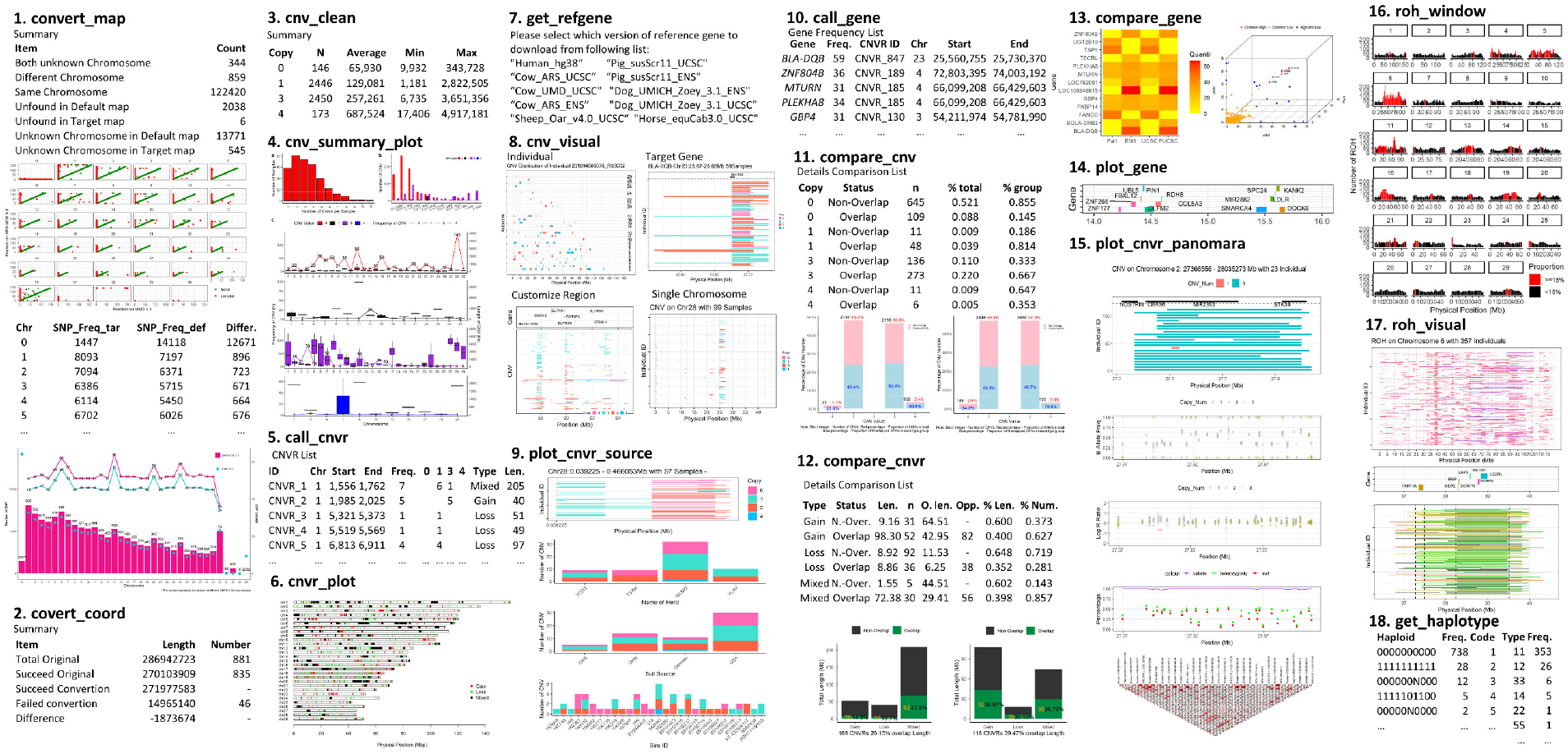
Main Functions and Work Flow.

The functions provided in this package can be categorised into five sections: Conversion; Summary; Annotation; Comparison; and Visualization. The conversion section is used to convert the SNP map files between different reference genomes and convert the coordinates for genomic intervals by given map files. The summary section is used to format, summarize, generate CNVR, make CNVR distribution maps, report high frequency ROH regions and generate haplotypes. The annotation section includes downloading and formatting of reference gene lists, and annotation of genes. The comparison section consists of comparison of CNV, comparison of CNVR, comparison of annotated gene frequency lists and the comparison between other intervals. The visualization section mainly focuses on customizing and integrating the plotting of all information related to CNV, ROH and high frequency CNVR. It supports making plots by chromosomes, individuals, interesting region or target gene, and can also customize the addition of the source, SNP genotype, signal intensity or linkage disequilibrium information.

## 3. Implementation

The package was developed with R Version 3.5.2, it was tested under Windows, OS and Linux systems. The detailed manual and vignette can be found at https://github.com/JH-Zhou/HandyCNV.

## Supporting information

Demo data and example

## Acknowledgements

This package depends on several developed R packages, such as Tidyverse family^1^ and data.table^2^. We appreciate all related contributors to the open source R language.

## Funding information

This project was funded by the China Scholarship Council.

## Notes

https://jh-zhou.github.io/HandyCNV/articles/HandyCNV.html

### Competing Interest Statement

The authors have declared no competing interest.

