## Supplementary material for "HandyCNV: Standardized Summary, Annotation, Comparison, and Visualization of CNV, CNVR and ROH": Demo data and example: HandyCNV_supplementary information.html

HandyCNV Vignette


### HandyCNV Vignette

###### February 14, 2021

- Introduction
  - Installation and Prerequisites
  - Demo data
  - Setting up a working directory
- What issues can this package solve?
  - 1. How do we prepare the standard cnv input file for `HandyCNV` and get a quick summary?
  - 2 How do we visualise CNVs?
    - 2.1 How do we have a quick overview of all CNVs on each chromosome?
    - 2.2 How do we plot CNVs in a region of interest?
    - 2.3 How do we plot CNVs in a sample of interest?
    - 2.4 How do we plot an interesting gene to see how many samples had a CNV overlapping with it?
    - 2.5 How can we present annotated genes above the CNV plot?
  - 3. What types of summary plots are available?
  - 4. How do we generate CNVRs (CNV Regions) from CNV results?
  - 5. How do we annotate genes for CNVRs and CNVs?
    - 5.1 How do we download the reference gene lists we need?
    - 5.2 The `call_gene` function
  - 6. How to plot CNVR distribution map?
  - 7. How can we plot all high frequency CNVRs at once?
  - 8. Can we compare CNVs between different result sets?
  - 9. What about CNVRs?
  - 10. How do we find the consensus set of genes common to multiple CNV result sets?
  - Brief summary
  - 11. How can we create the map file used in `HandyCNV`, to allow comparing CNVs between different reference genomes?
  - 12. How can we integrate the CNVs and annotated genes with additional information, such as Log R ratio, B Allele Frequency, call rate, heterozygosity, missing value rate and Linkage Disequilibrium, and plot it as one figure?
  - 13. How can we make a plot to show the source of the CNVs?
  - 14. How can we plot just the genes in a specific region, to save as a seperate figure?
  - 15. How can we find regions with high frequencies of runs of homozygosity?
  - 16. How do we visualise ROHs?
    - 16.1 Plot ROH by Chromsome
    - 16.2 Plot ROH by single region
    - 16.3 How can we present annotated genes above the ROH plot?
    - 16.4 How do we plot the full length of ROH for all samples that had a ROH in a given region?
    - 16.5 How do we plot ROHs on an interested sample?
  - 17. How do we get haplotype for ROH region?
  - 18. How do we convert coordinates for CNV, CNVR, ROH, or any other intervals?

### Introduction

This package was originally designed for the Post-analysis of CNV results inferred from PennCNV and CNVPartition (GenomeStudio). However, it has now been expanded to accept input files in standard formats for a wider range of applications. Our motivation is to provide a standard, reproducible and time-saving pipeline for the post-analysis of CNVs and ROHs detected from SNP genotyping data for the majority of diploid Species. The functions provided in this package can be categorised into five sections: Conversion, Summary, Annotation, Comparison and Visualization. The most useful features provided are: integrating summarized results, generating lists of CNVR, annotating the results with known gene positions, plotting CNVR distribution maps, and producing customised visualisations of CNVs and ROHs with gene and other related information on one plot. This package also supports a range of customisations, including the colour, size of high resolution figures, and output folder, avoiding conflict between the results of different runs. Running through all functions detailed in the vignette could help us to identify and explore the most interesting genomic regions more easily. In the following sections, we will present how to use the functions provided in this package to solve these problems.

#### Installation and Prerequisites

First, to run this package, we need to make sure that R (Version >= 3.5.2) is installed in your computer (R download link: https://www.r-project.org/). Once R is installed, the ‘HandyCNV’ package can be installed from Github repository by running the following script. If you rarely used R, it may take more time to install the ‘HandyCNV’ for the first time.

```
#install.packages("remotes")
#library(remotes)
#install_github(repo = "JH-Zhou/HandyCNV")
```

Then, we need to load the ‘HandyCNV’ package in order to run the following examples. This can be done using the `library` function as shown below.

```
library(HandyCNV)
```

To start playing with this package, we first need to prepare at least one CNV result list. With only CNV results as an input file, we can explore the functions from section 1 to section 10 as below. But to get more interesting and potentially valuable results, we will have to prepare additional input files, including the map file and signal intensity file for the SNP chip used to generate the CNV list, the pedigree of the samples, and Plink format (bed, bim and fam) genotype files. Many of these files are already required during CNV discovery, and by generating them early in the pipeline, you will find that the rest of workflow will be much easier and faster. We will be introducing how to prepare each of the input files in the relevant chunks below.

#### Demo data

We have provided some internal demo data, which should installed with package, in order to demonstrate how to use this package. We can access the demo data using the built-in R function `system.file` (see examples below). In the demo data, the first CNV results file is the default output from PennCNV with the ARS-UCD 1.2 (ARS) cattle reference genome. The second CNV results file contains the default output from CNVPartition with UMD 3.1 (UMD) cattle reference genome, and the third CNV list is an example of a standard input file that could be prepared from CNV results inferred by other tools.

Note: If you prefer not to test with this demo data, just skip this step to section 1 and use your own data instead.

```
cnv_penn_ars <- system.file("extdata", "Demo_data/cnv_results/ARS_PennCNV_WGC.goodcnv", package = "HandyCNV")
cnv_part_umd <- system.file("extdata", "Demo_data/cnv_results/UMD_CNVPartition.txt", package = "HandyCNV")
cnv_other_umd <- system.file("extdata", "Demo_data/cnv_results/UMD_Standard_CNV.txt", package = "HandyCNV")
```

#### Setting up a working directory

To start the analysis, let’s first set up the working directory. This will help ensure that all results will be saved in the same directory.

```
setwd(dir = "C:/Users/handy_test") #remember to replace the path with your own
```

### What issues can this package solve?

As above we mentioned, the functions in the package have been categorised into five sections: Conversion, Summary, Annotation, Comparison and Visualization.

Functions in the Conversion section can, for example, be used to convert coordinates between the default and target mapfiles provided by users; this would be useful if we are detecting CNVs with two different reference genomes. The converted map files can be output in formats suitable for use in Plink, PennCNV and our package ‘HandyCNV’. It is also possible to convert coordinates for intervals (CNV, CNVR, QTL et al) between the defaults and target maps, which might useful when comparing the results to those of other researchers. Functions included in the Summarization section allow the formatting and plotting of outputs, such as producing detailed summaries of CNVs and ROHs, making CNV summary plots, generating CNV regions (CNVRs), and plotting CNVR distribution maps.

The Annotation section allows the downloading of reference gene lists from the UCSC website, enabling the annotation of CNV, CNVR, ROH or other intervals with gene locations. The Comparison section facilitates the comparison of CNVs, CNVRs, Gene Frequent Lists and any other intervals. The highlight of this section is that comparison of CNVs and CNVRs will produce reports detailing comparison results on both individual and population levels, and all comparison results are relative to both input files. Finally, the Visualization section contains functions to support customised CNV and ROH plotting by chromosome, specific sample, regions of interest, or target genes (plotting by gene is not yet available for ROH). It is also possible to annotate the plots with other information, such as gene locations, log R ratio, B Allele Frequency, SNP genotype, LD or the source of CNVs. These functions can also plot genes separately from reference gene lists, which might useful for comparing outputs with plots from other studies.

Now, let’s start exploring the functions in `HandyCNV`.

#### 1. How do we prepare the standard cnv input file for `HandyCNV` and get a quick summary?

We can use `cnv_clean` function to convert cnv results to standard format. This function will also produce a brief summary of CNV results. First, let’s format the sample CNV results from PennCNV. Note:

1. When formatting results in PennCNV format, the corresponding CNV input argument is ‘penncnv’.
2. Default PennCNV results in Sample ID columns usually have the file path ahead the Sample ID. For example with the file “/home/zjhang027/C/final\_403\_penncnv/201094560082\_R03C01.txt”, the pattern ‘cnv/’ before the sample ID is unique, so we can take it as the separator using the argument `penn_id_sep = "cnv/"` to extract the wanted sample ID.

```
cnv <- cnv_clean(penncnv = cnv_penn_ars, #when running your own data, set this argument to 'penncnv = "localpath/file"'
                 penn_id_sep = "cnv/", #take the unique pattern as a separator to extract the sample ID 
                 drop_length = 5, # the maximum CNV length threshold (in Mb); CNVs larger than this value will be deleted
                 folder = "cnv_clean") #create a new folder to save results 
#> There are 382 individuals with 5215 CNVs in total.
#> The average number of CNVs present per individual is 13.65
#> Basic summary stats by CNV type:
#> # A tibble: 4 x 5
#>   CNV_Value     N `Average Length` `Min Length` `Max Length`
#>   <chr>     <int>            <dbl>        <dbl>        <dbl>
#> 1 0           146           65930.         9932       343728
#> 2 1          2446          129081.         1181      2822505
#> 3 3          2450          257261.         6735      3651356
#> 4 4           173          687524.        17406      4917181
```

Next, let’s format a second set of CNV results, this time produced using CNVPartition. Note: The corresponding input argument is `cnvpartition` for CNV result from CNVPartition.

```
cnv <- cnv_clean(cnvpartition = cnv_part_umd, #when running your own data, set this argument by 'cnvpartition = "localpath/file"' 
                 drop_length = 5, #the maximum CNV length threshold (in Mb); CNVs larger than this value will be deleted
                 folder = "cnv_clean") 
#> There are 365 individuals with 1239 CNVs in total.
#> The average number of CNV on each Individual is 3.39
#> Basic summary stats by CNV type:
#> # A tibble: 4 x 5
#>   CNV_Value     N `Average Length` `Min Length` `Max Length`
#>       <int> <int>            <dbl>        <dbl>        <dbl>
#> 1         0   754          148699.         4510      1204452
#> 2         1    59          375120.        14229      3440849
#> 3         3   409          840778.         4908      4867751
#> 4         4    17          634036.        58524      2336934
```

If the CNV results were not produced by PennCNV or CNVPartition, we need to arrange the CNV results into the standard format, then use the `standard_cnv` argument instead. The standard CNV input list should include the following columns (in order): `Sample_ID`, `Chr`, `Start`, `End` and `CNV_Value`.

```
tabl <- "
|Sample_ID           |Chr |Start   |End     |CNV_Value |
|--------------------|----|--------|--------|----------|
|201977910077_R05C01 |6   |9087392 |9250902 |1         |
|201094560084_R06C02 |7   |28583247|28621254|3         |
|201094560084_R06C02 |7   |37214745|37238472|1         |
|201094560067_R08C02 |4   |6248795 |6425277 |3         |
|201094560067_R08C02 |4   |66567442|66830563|3         |"
```

Let’s now run `cnv_clean` on the third Standard CNV input file. Note that we are using the argument `standard_cnv` for the third CNV result.

```
cnv <- cnv_clean(standard_cnv = cnv_other_umd, #when running your own data, set this argument by 'standard_cnv = "localpath/file"'
                 drop_length = 5, #the maximum CNV length threshold (in Mb); CNVs larger than this value will be deleted
                 folder = "cnv_clean")
#> There are 382 individuals with 4534 CNVs in total.
#> The average number of CNV per individual is 11.87
#> Basic summary stats by CNV type:
#> # A tibble: 4 x 5
#>   CNV_Value     N `Average Length` `Min Length` `Max Length`
#>       <int> <int>            <dbl>        <dbl>        <dbl>
#> 1         0    77           60964.         9932       343900
#> 2         1  2187          138295.            1      2843668
#> 3         3  2115          258623.            1      3933681
#> 4         4   155          779553.        17392      4869255
```

#### 2 How do we visualise CNVs?

The `cnv_visual` function supports visualising CNVs by chromosome, single sample, regions of interest, and target genes.

##### 2.1 How do we have a quick overview of all CNVs on each chromosome?

Once we got the standard `clean_cnv` result, we can use the `cnv_visual` function to plot all CNVs, to have a quick look if there are any notable patterns in CNV results.

First, let’s visualise the whole PennCNV dataset across all chromosomes:

```
cnv_visual(clean_cnv = "cnv_clean/penncnv_clean.cnv", #standard file was generated by 'cnv_clean' function in section 1
           max_chr = 29, #select how many chromosomes to plot 
           width_1 = 20, #optional,adjust the width of final plot
           height_1 =12, #optional,adjust the height of final plot
           folder = "cnv_visual_all_penn")
#> [1] "Input data passed requirment check..."
#> [1] "Plotting the CNVs on Chromosome 1"
#> [1] "Plotting the CNVs on Chromosome 2"
#> [1] "Plotting the CNVs on Chromosome 3"
#> [1] "Plotting the CNVs on Chromosome 4"
#> [1] "Plotting the CNVs on Chromosome 5"
#> [1] "Plotting the CNVs on Chromosome 6"
#> [1] "Plotting the CNVs on Chromosome 7"
#> [1] "Plotting the CNVs on Chromosome 8"
#> [1] "Plotting the CNVs on Chromosome 9"
#> [1] "Plotting the CNVs on Chromosome 10"
#> [1] "Plotting the CNVs on Chromosome 11"
#> [1] "Plotting the CNVs on Chromosome 12"
#> [1] "Plotting the CNVs on Chromosome 13"
#> [1] "Plotting the CNVs on Chromosome 14"
#> [1] "Plotting the CNVs on Chromosome 15"
#> [1] "Plotting the CNVs on Chromosome 16"
#> [1] "Plotting the CNVs on Chromosome 17"
#> [1] "Plotting the CNVs on Chromosome 18"
#> [1] "Plotting the CNVs on Chromosome 19"
#> [1] "Plotting the CNVs on Chromosome 20"
#> [1] "Plotting the CNVs on Chromosome 21"
#> [1] "Plotting the CNVs on Chromosome 22"
#> [1] "Plotting the CNVs on Chromosome 23"
#> [1] "Plotting the CNVs on Chromosome 24"
#> [1] "Plotting the CNVs on Chromosome 25"
#> [1] "Plotting the CNVs on Chromosome 26"
#> [1] "Plotting the CNVs on Chromosome 27"
#> [1] "Plotting the CNVs on Chromosome 28"
#> [1] "Plotting the CNVs on Chromosome 29"
#> [1] "Task done"
```

Figure.1 CNVs on Chromosome 15 (ARS reference) from PennCNV

Let’s now visualise CNVs from the CNVPartition results by chromosome. We can adjust the size of final figure by setting ‘width\_1’ and ‘height\_1’.

```
cnv_visual(clean_cnv = "cnv_clean/cnvpart_clean.cnv", #generated by 'cnv_clean' function in section 1
           max_chr = 29, #select how many Chromosomes to plot
           width_1 = 20, #optional,adjust the width of final plot
           height_1 = 12, #optional,adjust the height of final plot
           folder = "cnv_visual_all_part", )
```

Figure.2 CNVs on Chromosome 28 (UMD reference) from CNVPartition

##### 2.2 How do we plot CNVs in a region of interest?

After plotting all CNVs on each chromosome, we might find that some regions are particularly interesting. To plot these, we can add the options `chr_id`, `start_position` and `end_position` to the `cnv_visual` function call. Here, we are taking the region of Chr15:77-80.1221 Mb from the PennCNV results as an example.

```
cnv_visual(clean_cnv = "cnv_clean/penncnv_clean.cnv", #standard file was generated by 'cnv_clean' function in section 1
           chr_id = 15, #set Chromosome
           start_position = 77, #set start position, unit is 'Mb'
           end_position = 80.1221, #set end position, unit is 'Mb'
           col_0 = "red", #optional, customize colour for 0 copy CNVs 
           col_1 = "orange", #optional, customize colour for 1 copy CNVs 
           col_3 = "blue", #optional, customize colour for 3 copy CNVs 
           col_4 = "green", #optional, customize colour for 4 copy CNVs 
           width_1 = 13, #optional, adjust the width of final plot
           height_1 = 10, #optional, adjust the height of final plot
           folder = "cnv_visual")
#> [1] "Input data passed requirment check..."
#> [1] "Task done, plot was stored in working directory."
```

Figure.3 CNVs on Chr15:77-80.1221 Mb

##### 2.3 How do we plot CNVs in a sample of interest?

Here we need to assign both the Sample ID (to `individual_id`) and the clean CNV list (to `clean_cnv`) in the `cnv_visual` function. The individual CNV plot will saved in the current working directory. For example, the sample “201094560076\_R03C02” has the most CNVs in the PennCNV results, so let’s plot it and have a look. Note: we can use the `individual_cnv_count.txt` file (to be generated in section 3) to identify samples that exhibit large numbers of CNVs.

```
cnv_visual(clean_cnv = "cnv_clean/penncnv_clean.cnv", #standard file was generated by 'cnv_clean' function in section 1
           individual_id = "201094560076_R03C02", #set the sample ID  
           width_1 = 20, #optional, adjust the width of final plot
           height_1 = 13, #optional. adjust the height of final plot
           folder = "cnv_visual")
#> [1] "Input data passed requirment check..."
#> [1] "Task done, plot was stored in working directory."
```

Figure.4 CNVs in a specific sample

##### 2.4 How do we plot an interesting gene to see how many samples had a CNV overlapping with it?

The idea of plotting CNVs by target gene is to avoid making overly-strong statements when reporting important candidate genes after annotation for CNVRs. If we think that one particular gene is especially important in the CNV analysis, it’s better to present the frequency at which that gene has a structural variation in the experimental population. Plotting CNVs by gene will directly present how many samples show copy number variations overlapping the gene.

Note: The input file ‘cnv\_annotation.txt’ will be generated in section 5.2, as part of annotating the gene to CNV results.

```
cnv_visual(clean_cnv = "call_gene_Penn_UCSC/cnv_annotation.txt", #here we need the annotated CNV list, generated by 'call_gene' in section 5.2
           target_gene = "BLA-DQB", #set the gene name we are interested in
           col_0 = "darkorchid4",  #optional, customize colour for o copy CNV 
           col_1 = "dodgerblue3",  #optional, customize colour for 1 copy CNV 
           col_3 = "violetred3",  #optional, customize colour for 3 copy CNV 
           col_4 = "orangered3",  #optional, customize colour for 4 copy CNV
           width_1 = 13, #adjust the width of final plot
           height_1 = 10, #adjust the width of final plot
           folder = "cnv_visual")
#> [1] "Renamed 'CNV_Start' and 'CNV_End' as 'Start' and 'End', input data passed requirment check..."
#> [1] "Task done, plot was stored in working directory."
```

Figure.5 CNVs on a specific gene (*BLA-DQB*)

##### 2.5 How can we present annotated genes above the CNV plot?

To do this, we need to provide two input files to the `cnv_visual` function: the CNV list in `clean_cnv` and gene annotations in `refgene`. We should also set the genomic region we are interested in. The reference gene list will be generated by the `get_refgene` function in section 5.

```
cnv_visual(clean_cnv = "cnv_clean/penncnv_clean.cnv", #standard file was generated by 'cnv_clean' function in section 1
           refgene = "refgene/cow_ARS_UCSC.txt", #standard input file will generated by the function 'get_refgene' in Section 5
           chr_id = 23, #set Chromosome
           start_position = 25.56, #set start position, unit is 'Mb'
           end_position = 25.74, #set end position, unit is 'Mb'
           gene_font_size = 2.2, #optional, adjust the size of gene font
           show_name = c(25.66,27.72), #optional, show the gene in the given interval, maximum support two intervals
           col_0 = "tomato", #optional, customize colour for 0 copy CNV 
           col_1 = "hotpink", #optional, customize colour for 1 copy CNV 
           col_3 = "turquoise", #optional, customize colour for 3 copy CNV 
           col_4 = "aquamarine", #optional, customize colour for 4 copy CNV 
           col_gene = "red", #optional, customize colour of gene  
           width_1 = 14, #optional, adjust the width of final plot
           height_1 = 10, #optional, adjust the height of final plot
           folder = "cnv_visual")
#> [1] "Input data passed requirment check..."
#> [1] "CNV on Chromosome 23: 25.56 - 25.74 Mb with 68 Individual - "
#> [1] "Plotting gene..."
#> [1] "Task done, plot was stored in working directory."
```

Figure.6 CNVs with annotated genes

#### 3. What types of summary plots are available?

Using the clean CNV results, we can run the `cnv_summary_plot` function to generate a range of summary plots. These plots can aggregate CNV results by length group, CNV type, chromosome, and individual. It is also possible to make two different types of plots combining these figures.

```
cnv_summary_plot(clean_cnv = "cnv_clean/cnvpart_clean.cnv", #standard file was generated by 'cnv_clean' function in section 1
                 length_group = c(0.05, 0.1, 0.2, 0.3, 0.4, 0.5, 0.6, 0.7, 1, 2, 5), # optional, set group of vectors to divide CNV length, unit is Mb. such as vector of ‘c(0.05, 0.3, 1)’, means divide the CNV length into four group: '<0.05Mb', '0.05 - 0.3Mb', '0.3-1Mb' and '>1Mb', maximum can accept 11 values
                 col_0 = "red", #optional, customize color for o copy CNV 
                 col_1 = "black", #optional, customize color for o copy CNV 
                 col_3 = "purple", #optional, customize color for o copy CNV 
                 col_4 = "blue", #optional, customize color for o copy CNV 
                 plot_sum_1 = TRUE,  #optional, make sum combination plot 1
                 height_sum1 = 26, #optional, adjust the size of sum plot 1
                 width_sum1 = 20, #optional, adjust the size of sum plot 1
                 plot_sum_2 = TRUE, #optional, make sum combination plot 2
                 height_sum2 = 20, #optional, adjust the size of sum plot 2
                 width_sum2 = 27, #optional, adjust the size of sum plot 2
                 folder = "cnv_summary_plot_penn")
#> The CNV distribution plot has been saved in the 'cnv_summary_plot_penn' directory.
#> The CNV length group plot was saved in the 'cnv_summary_plot_penn' directory.
#> The box plot of CNV lengths was saved in the 'cnv_summary_plot_penn' directory.
#> [1] "Distribution of CNV type on chromosome plot was saved in working directory."
#> [1] "The number of CNV in Individual plot was saved in working directory."
#> [1] "cnv_summary_plot_1 was saved in working directory."
#> [1] "cnv_summaray_polt_2 was saved in working directory."
#> [1] "Task done."
```

Figure.7 CNV Summary Plot Combination 1

Figure.8 CNV Summary Plot Combination 2

#### 4. How do we generate CNVRs (CNV Regions) from CNV results?

CNV regions are defined as regions of the genome containing one or more CNVs that overlap by at least one base pair: s a result, CNVRs will never overlap one another. As an example, let’s first generate CNVRs for the clean PennCNV results, using the `call_cnvr` function. A key parameter of this function is `chr_set`, which allows setting the maximum number of chromosomes over which to generate CNVRs. In our example, the species is *Bos taurus* (cows), with 29 autosomes, so we set `chr_set = 29`. This function will output three tables:

1. The list of CNVRs. This is saved into the file `individual_cnv_cnvr.txt`.
2. A brief summary table showing numbers of CNVRs by length and type (Deletion, Duplication, and Mixed). Mixed indicates that both duplications and deletions are found within the CNVR.
3. The total length and number of CNVRs on each chromosome.

The resulting CNVRs are sorted by chromosome, start position, and end position, then assigned the CNVR ID to each unique interval by order. One of the output files, `individual_cnv_cnvr.txt` contain both CNVR and CNV information for every sample: this file might suitable for CNV-GWAS analyses.

```
cnvr <- call_cnvr(clean_cnv = "cnv_clean/penncnv_clean.cnv", #standard file was generated by 'cnv_clean' function in section 1
                  chr_set = 29, #Set the maximum number of chromosomes (29 autosomes for cattle)
                  folder = "call_cnvr_penn")
#> [1] "Chromosome 1 has been processed."
#> [1] "Chromosome 2 has been processed."
#> [1] "Chromosome 3 has been processed."
#> [1] "Chromosome 4 has been processed."
#> [1] "Chromosome 5 has been processed."
#> [1] "Chromosome 6 has been processed."
#> [1] "Chromosome 7 has been processed."
#> [1] "Chromosome 8 has been processed."
#> [1] "Chromosome 9 has been processed."
#> [1] "Chromosome 10 has been processed."
#> [1] "Chromosome 11 has been processed."
#> [1] "Chromosome 12 has been processed."
#> [1] "Chromosome 13 has been processed."
#> [1] "Chromosome 14 has been processed."
#> [1] "Chromosome 15 has been processed."
#> [1] "Chromosome 16 has been processed."
#> [1] "Chromosome 17 has been processed."
#> [1] "Chromosome 18 has been processed."
#> [1] "Chromosome 19 has been processed."
#> [1] "Chromosome 20 has been processed."
#> [1] "Chromosome 21 has been processed."
#> [1] "Chromosome 22 has been processed."
#> [1] "Chromosome 23 has been processed."
#> [1] "Chromosome 24 has been processed."
#> [1] "Chromosome 25 has been processed."
#> [1] "Chromosome 26 has been processed."
#> [1] "Chromosome 27 has been processed."
#> [1] "Chromosome 28 has been processed."
#> [1] "Chromosome 29 has been processed."
#> 978 CNVRs generated in total.
#> Overall summary of CNVRs:
#> # A tibble: 3 x 6
#>   Type      N `Average Length` `Min Length` `Max Length` `Total Length`
#>   <chr> <int>            <dbl>        <dbl>        <dbl>          <dbl>
#> 1 Gain    347          150153.         7947      2639041       52103038
#> 2 Loss    399          100664.         3703       737404       40164889
#> 3 Mixed   232          898204.         6735      8282577      208383407
#> Summary of CNVRs on each Chromosome:
#> # A tibble: 29 x 3
#>      Chr `Total Length` `Number of CNVR`
#>    <int>          <dbl>            <int>
#>  1     1       26386484               62
#>  2     2       18333125               51
#>  3     3       11339395               39
#>  4     4       15529397               56
#>  5     5       12595686               53
#>  6     6       28976727               44
#>  7     7       14807963               52
#>  8     8       10681061               24
#>  9     9       14508332               37
#> 10    10        6784489               35
#> # ... with 19 more rows
```

Next, let’s generate CNVRs for CNVPartition and standard CNV lists. In order to avoid mixing the results among different files, we can set the `folder` argument to assign a distinguishing name to the newly generated folder.

```
call_cnvr(clean_cnv = "cnv_clean/cnvpart_clean.cnv", #standard file was generated by 'cnv_clean' function in section 1
                    chr_set = '29', 
                    folder = "call_cnvr_part")

call_cnvr(clean_cnv = "cnv_clean/cleancnv.cnv", #standard file was generated by 'cnv_clean' function in section 1
          chr_set = 29,
          folder = "call_cnvr_standard")
```

#### 5. How do we annotate genes for CNVRs and CNVs?

Once we have defined some CNV regions, we might want to see how many genes are located each. We can annotate genes to both CNVs and CNVRs using the `call_gene` function. Before that, however, we need to prepare gene lists for the correct reference genome. This can be done using the `get_refgene` function, as described in the following sections.

##### 5.1 How do we download the reference gene lists we need?

We can use the `get_refgene` function to automatically download the reference gene list and convert it to the standard format required for functions such as `call_gene`. First, let’s run `get_refgene` without arguments to see which reference gene lists we can download. If you don’t find the information you’re looking for, please feel free to contact our maintainer to update it to you. For now, only reference gene lists from the UCSC website are supported; however, we may add more features if there is sufficient demand.

```
get_refgene()
#> Please select which version to download from following list:
#>  [1] "Cow_ARS_UCSC"            "Cow_ARS_ENS"            
#>  [3] "Cow_UMD_UCSC"            "Pig_susScr11_UCSC"      
#>  [5] "Pig_susScr11_ENS"        "Human_hg38"             
#>  [7] "Sheep_Oar_v4.0_UCSC"     "Horse_equCab3.0_UCSC"   
#>  [9] "Dog_UMICH_Zoey_3.1_UCSC" "Dog_UMICH_Zoey_3.1_ENS"
```

Now, let’s get the reference gene list for the PennCNV results. This dataset was generated using ARS-UCD 1.2 as reference genome. Two gene lists are available for this reference: `cow_ARS_UCSC` and `cow_ARS_ENS`. The suffix `ENS` indicates the Ensembl reference gene list. The gene list will be saved in the folder `refgene`, which will be created first if necessary.

```
get_refgene(gene_version = "Cow_ARS_UCSC")
#> [1] "Link to the website..."
#> [1] "Converting format..."
#> [1] "Task done."
get_refgene(gene_version = "Cow_ARS_ENS")
#> [1] "Link to the website..."
#> [1] "Converting format..."
#> [1] "Task done."
```

Next let’s get reference gene list for the CNVPartition and Standard result sets, both of which were generated on the UMD 3.1 reference genome; in this case, `Cow_UMD_UCSC` is the correct option to select.

```
get_refgene(gene_version = "Cow_UMD_UCSC")
#> [1] "Link to the website..."
#> [1] "Converting format..."
#> [1] "Task done."
```

Now we have well-prepared reference gene lists, let’s return to the `call_gene` function.

##### 5.2 The `call_gene` function

The `call_gene` function has three arguments:

1. `ref_gene`: this argument is required, and takes the path of a downloaded gene set (in the `refgene` folder).
2. `interval`: an optional file containing a list of CNV regions. This argument can also point to any file storing interval data in the correct format; examples could include CRVs, QTL windows, or haplotype lists.
3. `clean_cnv`: an optional file containing a list of CNVs

We suggest that both ‘interval’ and ‘clean\_cnv’ should be set simultaneously: this will enable reporting of the frequency of all annotated genes by counting how many CNVs overlapped to each gene within a CNVR. In section 10, we will show how to use the output file `gene_freq_cnv.txt` to make comparisons between different CNV results to find sets of consensus genes.

As an example, we will first call genes for the PennCNV results, ensuring that the gene list for the correct reference has been set (ARS 1.2). In this case, we are using the Ensembl-provided gene list.

```
call_gene(refgene = "refgene/cow_ARS_ENS.txt", #standard file was generated by 'get_refgene' function in section 5.1
          interval = "call_cnvr_Penn/cnvr.txt", #optional, standard file was generated by 'call_cnvr' function in section 4
          clean_cnv = "cnv_clean/penncnv_clean.cnv", #optional, standard file was generated by 'cnv_clean' function in section 1
          folder = "call_gene_Penn_Ens")
#> Checking gene annotation status in the interval file...
#> Summary of annotation results:
#>   Interval_Has_Gene Interval_Without_Gene Total_Number_of_Genes
#> 1               692                   286                  2906
#> Checking gene annotation status in CNV file...
#> 2906 genes were matched in the CNV and CNVR results. The 10 most frequent genes:
#> # A tibble: 10 x 2
#> # Groups:   name2 [10]
#>    name2   Frequency
#>    <chr>       <int>
#>  1 53092.3        72
#>  2 19939.4        67
#>  3 49535.4        67
#>  4 77856.1        63
#>  5 08346.6        60
#>  6 31707.5        59
#>  7 BLA-DQB        59
#>  8 75223.1        51
#>  9 33864.5        48
#> 10 53751.3        48
#> Warning in call_gene(refgene = "refgene/cow_ARS_ENS.txt", interval =
#> "call_cnvr_Penn/cnvr.txt", : The following genes are duplicated in multiple
#> CNVRs, and will only be annotated on the first CNVR_ID in the final gene
#> frequency list report!
#>            name2       ID Chr   i.Start     i.End
#> 49            U7  CNVR_35   1  47249572  49067075
#> 233       ZNF644 CNVR_128   3  52257274  52264697
#> 234       ZNF644 CNVR_129   3  52321114  52363402
#> 315         DGKB CNVR_166   4  22417069  22677610
#> 316         DGKB CNVR_167   4  23108150  23239528
#> 361        KCND2 CNVR_195   4  84792163  84829041
#> 362        KCND2 CNVR_196   4  84948868  85008727
#> 494      84617.1 CNVR_258   5 118326410 118372340
#> 495        TAFA5 CNVR_258   5 118326410 118372340
#> 496      84617.1 CNVR_259   5 118421130 118474468
#> 497        TAFA5 CNVR_259   5 118421130 118474468
#> 520           U6 CNVR_272   6  30915027  33530543
#> 521       CCSER1 CNVR_272   6  30915027  33530543
#> 522       CCSER1 CNVR_273   6  33709997  33774948
#> 523       CCSER1 CNVR_274   6  33873388  34610791
#> 859           U2 CNVR_349   7  84759907  86193204
#> 863          7SK CNVR_352   7  91189501  92330502
#> 915  Metazoa_SRP CNVR_384   9   1492214   5583983
#> 931        GRIK2 CNVR_396   9  45904339  47944789
#> 932        GRIK2 CNVR_397   9  48307662  48796022
#> 967      5S_rRNA CNVR_424  10   9143166   9354675
#> 1059 Metazoa_SRP CNVR_439  10  42661079  42860796
#> 1064      UNC13C CNVR_442  10  55628092  55693450
#> 1065      UNC13C CNVR_443  10  55907151  55929569
#> 1074         7SK CNVR_449  10  94536813  96569247
#> 1104          U7 CNVR_461  11  17249565  18200634
#> 1119 Metazoa_SRP CNVR_471  11  51936197  54294891
#> 1300     5S_rRNA CNVR_498  12  18874232  18999163
#> 1660     5S_rRNA CNVR_611  15  69398047  70655610
#> 1735     5S_rRNA CNVR_628  16  16011331  16812729
#> 1824        LRBA CNVR_651  17   6794956   6825613
#> 1825        LRBA CNVR_652  17   6963919   7013884
#> 1972        WWOX CNVR_682  18   5512831   5529148
#> 1973        WWOX CNVR_683  18   5549190   5640099
#> 2005       ITFG1 CNVR_692  18  15578052  15614150
#> 2006       ITFG1 CNVR_693  18  15714407  15821257
#> 2160       ACACA CNVR_728  19  13518465  13564941
#> 2161       ACACA CNVR_729  19  13678370  13749514
#> 2797        BPHL CNVR_859  23  50437086  50536890
#> 2798        BPHL CNVR_860  23  50546856  50644819
#> 2809          U2 CNVR_867  24  10709058  10979040
#> 3022     5S_rRNA CNVR_900  25  41532834  41632501
#> 3055      ATRNL1 CNVR_912  26  35555953  35700362
#> 3056      ATRNL1 CNVR_913  26  35829748  35904911
#> 3104      JMJD1C CNVR_938  28  18916753  19430612
#> 3105      JMJD1C CNVR_939  28  19507984  19563947
#> 3108      CTNNA3 CNVR_941  28  20997159  22372439
#> 3109      CTNNA3 CNVR_942  28  22771955  22933760
#> 3110      CTNNA3 CNVR_943  28  23532241  23555163
#> 3112      CTNNA3 CNVR_944  28  24001450  24095133
#> 3196          U6 CNVR_975  29  48495358  48602322
#> Task done. Files containing the CNVs, CNVR annotations, summary tables, and the frequency of genes in CNV regions have been saved in the 'call_gene_Penn_Ens'.
```

We can also use the UCSC gene list:

```
call_gene(refgene = "refgene/cow_ARS_UCSC.txt", #standard file was generated by 'get_refgene' function in section 5.1
          interval = "call_cnvr_Penn/cnvr.txt", #standard file was generated by 'call_cnvr' function in section 4
          clean_cnv = "cnv_clean/penncnv_clean.cnv", #optional, standard file was generated by 'cnv_clean' function in section 1
          folder = "call_gene_Penn_UCSC")
#> Warning in call_gene(refgene = "refgene/cow_ARS_UCSC.txt", interval =
#> "call_cnvr_Penn/cnvr.txt", : The following genes are duplicated in multiple
#> CNVRs, and will only be annotated on the first CNVR_ID in the final gene
#> frequency list report!
```

Comparing the results, we observe that the results from two reference gene lists are slightly different, likely because the maintainers of each database have different methodologies (and possibly data sources) for producing their gene annotations. As a result of this, the position and quantity of genes might slightly differ between gene builds even for the same reference genome. Therefore, it may be important to validate genes of interest in more than one database.

Now, let’s call genes for the CNVPartition results. As before, we provide both CNVR and CNV lists, and set the new folder name by arguments. Note that we need to use the UMD 3.1 reference build for this data set. We also run the standard CNV set (using the ARS1.2 reference).

```
call_gene(refgene = "refgene/cow_UMD_UCSC.txt", #standard file was generated by 'get_refgene' function in section 5.1
          interval = "call_cnvr_Part/cnvr.txt", #standard file was generated by 'call_cnvr' function in section 4
          clean_cnv = "cnv_clean/cnvpart_clean.cnv", #optional, standard file was generated by 'cnv_clean' function in section 1
          folder = "call_gene_Part")
#> Warning in call_gene(refgene = "refgene/cow_UMD_UCSC.txt", interval =
#> "call_cnvr_Part/cnvr.txt", : The following genes are duplicated in multiple
#> CNVRs, and will only be annotated on the first CNVR_ID in the final gene
#> frequency list report!

call_gene(refgene = "refgene/cow_ARS_UCSC.txt", #standard file was generated by 'get_refgene' function in section 5.1
          interval = "call_cnvr_standard/cnvr.txt", #standard file was generated by 'call_cnvr' function in section 4
          clean_cnv = "cnv_clean/cleancnv.cnv", #optional, standard file was generated by 'cnv_clean' function in section 1
          folder = "call_gene_standard_ARS_UCSC")
#> Warning in call_gene(refgene = "refgene/cow_ARS_UCSC.txt", interval =
#> "call_cnvr_standard/cnvr.txt", : The following genes are duplicated in multiple
#> CNVRs, and will only be annotated on the first CNVR_ID in the final gene
#> frequency list report!
```

#### 6. How to plot CNVR distribution map?

The function `cnvr_plot` is used to plot the CNVR distribution across chromosomes. For this function, we only need to assign the a CNVR list file to the `cnvr` argument, and the reference genome assembly to the `assembly` argument (to provide chromosome lengths). For Bovine data, we provide data for both ARS and UMD. For other species with data available (see the `ref_gene` function in section 5.1), we can just assign the species name to the `assembly` argument, for example, `assembly = "human"` for human data.

First, we’ll plot the CNVR distribution for the PennCNV data set. The default legend position might not suitable for all species; the arguments `legend_x` and `legend_y` can be used to move it if necessary.

```
cnvr_plot(cnvr = "call_cnvr_Penn/cnvr.txt", #standard file was generated by 'call_cnvr' function in section 4
          assembly = "ARS", #cattle could select 'UMD' or 'ARS', other species could assign the name of species
          loss_col = "green2", #optional, adjust the color of Loss type of CNVR
          gain_col = "red", #optional, adjust the color of Gain type of CNVR
          mixed_col = "black",  #optional, adjust the color of Mixed type of CNVR
          legend_x = 127, #optional, adjust the horizontal position of legend
          legend_y = 30,  #optional, adjust the vertical position of legend
          height_1 = 14, #optional, adjust the height of CNVR plot
          width_1 = 10, #optional, adjust the width of CNVR plot
          folder = "cnvr_plot_penn")
#> [1] "Task done, CNVR Distribution Plot saved in working directory."
```

Figure.9 CNVR distribution map for PennCRV data

Next, we plot the CNVRs derived from the CNVPartition data set.

```
cnvr_plot(cnvr = "call_cnvr_Part/cnvr.txt", #standard file was generated by 'call_cnvr' function in section 4
          assembly = "UMD", #cattle could select 'UMD' or 'ARS', other species could assign the name of species
          loss_col = "darkorchid", #optional, adjust the color of Loss type of CNVR
          gain_col = "gold", #optional, adjust the color of Gain type of CNVR
          mixed_col = "tomato",  #optional, adjust the color of Mixed type of CNVR
          legend_x = 127, #optional, adjust the horizontal position of legend
          legend_y = 30,  #optional, adjust the vertical position of legend
          height_1 = 14, #optional, adjust the height of CNVR plot
          width_1 = 10, #optional, adjust the width of CNVR plot
          folder = "cnvr_plot_part")
#> [1] "Task done, CNVR Distribution Plot saved in working directory."
```

Figure.10 Part CNVR Distribution Map

#### 7. How can we plot all high frequency CNVRs at once?

It always interesting to see which genomic regions have the highest frequencies of structure variation. To view these, we can plot all the high frequency regions using the `cnvr_plot` function.

As an example, let’s plot the PennCNV results. Remember to set a new folder name to save the results.

```
cnvr_plot(cnvr = "call_cnvr_Penn/cnvr.txt", #standard file was generated by 'call_cnvr' function in section 4
          clean_cnv = "cnv_clean/penncnv_clean.cnv", #standard file was generated by 'cnv_clean' function in section 1
          refgene = "refgene/cow_ARS_ENS.txt", #standard file was generated by 'get_refgene' function in section 5.1
          sample_size = 382, #total sample size
          common_cnv_threshold = 0.18, #the CNV frequency threshold
          width_1 = 10, #optional, adjust the width of final plot
          height_1 = 10, #optional, adjust the height of final plot
          col_gene = "blue", #optional, adjust the colour of annotated genes
          gene_font_size = 2.5, #optional, adjust the font size of annotated gene names
          folder = "high_freq_cnvr_penn")
```

Figure.11 All high frequency CNVRs from PennCNV

#### 8. Can we compare CNVs between different result sets?

Typically, we will use multiple models and software packages when predicting and analysing CNVs, especially when using data based on SNP chip genotypes. As a result, it is very common that we will need to compare two or more sets of predicted CNVs. To tackle this problem, we have implemented the `compare_cnv` function to make comparisons on both the individual and population levels. At the individual level, the function checks how many CNVs overlap in the same individual between two CNV result sets, thereby verifying the level of similarity between the two sets of results. On the population level, the comparison will check the level of overlap through the whole sample population, without checking whether or not the CNVs are present in the same individual; this can better indiciate the degree of repeatability of CNV detection between the two result sets.

As an example let’s compare the CNV prediction results for PennCNV and CNVPartition. Note that these results were generated using different reference builds, so the results of the comparison are currently rather meaningless (the physical coordinates of the CNVs will not line up across reference builds): this problem will be fixed in section 11 below.

```
compare_cnv(cnv_def = "cnv_clean/cleancnv.cnv", #standard file was generated by 'cnv_clean' function in section 1
            cnv_tar = "cnv_clean/penncnv_clean.cnv", #standard file was generated by 'cnv_clean' function in section 1
            width_1 = 12, #optional, adjust the width of final plot
            height_1 = 11, #optional, adjust the height of final plot
            legend_x = 0.9, #optional, adjust the horizontal position of final plot
            legend_y = 0.9, #optional, adjust the vertical position of final plot
            col_1 = "pink", #optional, adjust the colour of Non-Overlapped part
            col_2 = "lightblue", #optional, adjust the colour of Overlapped part
            plot_caption = TRUE, #optional, show the Note information in plot or not
            folder = "compare_cnv_PartUMD_Vs_PennUMD")
#> [1] "Comparison Passed Validation, the number of Overlap and Non-overlap CNV equal to the original number of CNVs which are in total 4534"
#> [1] "The number of overlaped CNVs on individual level is 951, which is around 21 percent in the first file"
#> [1] "The number of Non-overlaped CNVs on individual level is 3583, which is around 79 percent in the first file"
#> [1] "Comparison to the first file was Passed Validation, the number of Overlap and Non-overlap CNV equal to the original number of CNVs which are in total of 4534"
#> [1] "The number of overlaped CNVs on population level is 2352, which is around 51.9 percent in first file."
#> [1] "The number of Non-overlaped CNVs on population level is 2182, which is around 48.1 percent in first file"
#> [1] "The final comparison results of the first file as follows: "
#>                  Item Individual.Level Population.Level
#> 1     Overlapped CNVs               21             51.9
#> 2 Non-overlapped CNVs               79             48.1
#> [1] "Comparison to the first file was finished."
#> [1] "Comparison to second file was Passed Validation, the number of Overlap and Non-overlap CNV equal to the original number of CNVs which are in total 5215"
#> [1] "The number of overlaped CNVs on individual level is 966, which is around 18.5 percent in second file."
#> [1] "The number of Non-overlaped CNVs on individual level is 4249, which is around 81.5 percent in second file."
#> [1] "Comparison to the second file was Passed Validation, the number of Overlap and Non-overlap CNV equal to the original number of CNVs which are in total of 5215"
#> [1] "The number of overlaped CNVs in seceond file on population level are 2575, which is around 49.4 percent"
#> [1] "The number of Non-overlaped CNVs in second file on population level are 2640, which is around 50.6 percent"
#> [1] "The final comparison results of the second file as follows: "
#>                  Item Individual.Level Population.Level
#> 1     Overlapped CNVs             18.5             49.4
#> 2 Non-overlapped CNVs             81.5             50.6
#> [1] "Task done. Comparison results were saved in your working directory"
```

Figure.12 Comparison of PennCNV and CNVPartition results

#### 9. What about CNVRs?

Similar to the `compare_cnv` function, we can use the `compare_cnvr` function to compare CNV regions. The same caveats apply with regard to the ensuring the two CNVR data sets were built on the same reference genome. Let’s try to use `compare_cnvr` now with the CNVPartition and Standard data sets (as before, ignoring for now the fact that these were produced on different reference genomes).

```
compare_cnvr(cnvr_def  = "call_cnvr_Part/cnvr.txt", #standard file was generated by 'call_cnvr' function in section 4
             cnvr_tar  = "call_cnvr_standard/cnvr.txt", #standard file was generated by 'call_cnvr' function in section 4
             width_1 = 8, #optional, adjust the width of comparison plot
             height_1 = 8, #optional, adjust the height of comparison plot
             hjust_prop = 0.2, #optional, adjust the horizontal position of the proportion number present in plot
             hjust_num = 1.6, #optional, adjust the horizontal position of the number present in plot
             col_1 = "grey15", #optional, adjust the colour of non-overlapped part
             col_2 = "forestgreen", #optional, adjust the colour of overlapped part
             folder = "compare_cnvr_PartUMD_Penn_UMD")
#> Comparison to the first file was Passed Validation, the number of Overlap and Non-overlap CNV equal to the original number of CNVRs which are in total of 246 
#> CNVR comparison summary results in checkover_pop_1 as following:
#> # A tibble: 6 x 8
#> # Groups:   Type [3]
#>   Type  Check_overlap origi_length num_CNVR overlap_len num_overlap_opp~
#>   <chr> <chr>                <int>    <int>       <dbl>            <int>
#> 1 Gain  Non-Overlap        1824550        7    53469808                0
#> 2 Gain  Overlap          105638930       76    53993672              100
#> 3 Loss  Non-Overlap        6194700       64     9566899                0
#> 4 Loss  Overlap           11580677       64     8208478               64
#> 5 Mixed Non-Overlap        3317779        4    36001939                0
#> 6 Mixed Overlap           70606375       31    37922215               60
#> # ... with 2 more variables: prop_overlap_len <dbl>, prop_num <dbl>
#> The number of overlaped CNVRs on population level is 171, which is around 69.5 percent in first file.
#> The number of Non-overlaped CNVRs on population level is 75, which is around 30.5 percent in first file
#> The overlapping length is 100124365 bp, which is around 50.3 percent in first file
#> The final comparison results of the first file as follows: 
#>                   Item Number.of.CNVR Proportion.of.Number.... Length.bp.
#> 1     Overlapped CNVRs            171                     69.5  100124365
#> 2 Non-overlapped CNVRs             75                     30.5   99038646
#> 3             In Total            246                    100.0  199163011
#>   Proportion.of.Length....
#> 1                     50.3
#> 2                     49.7
#> 3                    100.0
#> Comparison to the first file was finished.
#> Comparison to the second file was Passed Validation, the number of Overlap and Non-overlap CNV equal to the original number of CNVRs which are in total of 878
#> CNVR comparison summary results in checkover_pop_2 as following:
#> # A tibble: 6 x 8
#> # Groups:   Type [3]
#>   Type  Check_overlap origi_length num_CNVR overlap_len num_overlap_opp~
#>   <chr> <chr>                <int>    <int>       <dbl>            <int>
#> 1 Gain  Non-Overlap       35340763      238    36131115                0
#> 2 Gain  Overlap           13315879       68    12525527               68
#> 3 Loss  Non-Overlap       31958106      311    32810816                0
#> 4 Loss  Overlap            6279030       51     5426320               51
#> 5 Mixed Non-Overlap       70399922      107   117876424                0
#> 6 Mixed Overlap          129649020      103    82172518              106
#> # ... with 2 more variables: prop_overlap_len <dbl>, prop_num <dbl>
#> The number of overlaped CNVRs in seceond file on population level are 222, which is around 25.3 percent
#> The number of Non-overlaped CNVRs in second file on population level are 656, which is around 74.7 percent
#> The overlapping length is 100124365 bp, which is around 34.9 percent in second file
#> The final comparison results of the second file as follows: 
#>                   Item Number.of.CNVR Proportion.of.Number.... Length.bp.
#> 1     Overlapped CNVRs            222                     25.3  100124365
#> 2 Non-overlapped CNVRs            656                     74.7  186818355
#> 3             In Total            878                    100.0  286942720
#>   Proportion.of.Length....
#> 1                     34.9
#> 2                     65.1
#> 3                    100.0
#> Processing the unique common CNVRs list by uniting overlepped CNVRs between two results 
#> There are 168 unique and mutual CNVRs after uniting the overlapped CNVRs between two results 
#> Task done. Comparison results were saved in the working directory
```

Figure.13 Comparison of CNVR

#### 10. How do we find the consensus set of genes common to multiple CNV result sets?

One of the most interesting parts of CNV analysis is to find out which genes are affected by structure variations, and to check out the frequency of these aberrant genes in our samples. After using the `call_gene` function in section 5.2, we now have the gene frequency list for each CNV result set. We can take these lists as input in the `compare_gene` function to find the set of consensus genes common to multiple CNV results.

Now, let’s compare the annotated gene lists for the PennCNV and CNVPartition results as an example. We can assign the arguments `gene_freq_1` and `gene_freq_2` with the `gene_freq_cnv.txt` files from the `call_gene_Part` and `call_gene_Penn` output folders created earlier. Then we need to set an integer threshold (`common_gene_threshold`) for the number of observations required to declare that a gene is “commom”. In our example, the sample size is 400, so for a prevalence of 5%, the `common_gene_threshold` could calculated as `400 * 5% = 20`.

```
compare_gene(gene_freq_1 = "call_gene_Part/gene_freq_cnv.txt", #standard file was generated by 'call_gene' function in Section 5.2
             gene_freq_2 = "call_gene_Penn_Ens/gene_freq_cnv.txt", #standard file was generated by 'call_gene' function in Section 5.2
             title_1 = "Bovine Part", #optional, adjust the title of input file 1 present in final plot 
             title_2 = "Bovine Penn", #optional, adjust the title of input file 2 present in final plot 
             common_gene_threshold = 20, #optional, the threshold is the gene frequency, here 20 means the Gene frequency exceed 20 in two input lists will be regarded as common gene. 
             col_1 = "red", #optional, set the colour of 'Common High' genes
             col_2 = "blue", #optional, set the colour of 'Common Low' genes
             col_3 = "gold", #optional, set the colour of 'gene_freq_1'-only genes
             col_4 = "green", #optional, set the colour of 'gene_freq_2'-only genes
             height_1 = 10, #optional, adjust the height of final plot
             width_1 =  12, #optional, adjust the width of final 
             folder = "compare_gene_Part_Vs_Penn")
#> [1] "Clustering common gene between the two input files..."
#> [1] "Brief summary of comparison as below:"
#> # A tibble: 4 x 2
#> # Groups:   Common_Gene [4]
#>   Common_Gene          n
#>   <chr>            <int>
#> 1 Common_High          5
#> 2 Common_Low        3035
#> 3 High_Freq_list_1     6
#> 4 High_Freq_list_2    54
#> [1] "Gene comparison list, brife summary and comparison plot was saved in working directory."
```

Figure.14 Comparison of genes present in the PennCNV (green) and CNVPartition (gold) gene lists

We can also compare the frequency lists for three genes, just by adding one more input file (using the `gene_freq_3` argument); in this case, the comparison figure will be a 3D plot.

```
compare_gene(gene_freq_1 = "call_gene_Part/gene_freq_cnv.txt", #standard file was generated by 'call_gene' function in Section 5.2
             gene_freq_2 = "call_gene_Penn_Ens/gene_freq_cnv.txt", #standard file was generated by 'call_gene' function in Section 5.2
             gene_freq_3 = "call_gene_standard_ARS_UCSC/gene_freq_cnv.txt", #standard file was generated by 'call_gene' function in Section 5.2
             title_1 = "Part", #optional, adjust the title of input file 1 present in final plot 
             title_2 = "ENS", #optional, adjust the title of input file 2 present in final plot 
             title_3 = "UCSC", #optional, adjust the title of input file 3 present in final plot 
             common_gene_threshold = 15,
             height_1 = 5.8, #optional, adjust the height of final plot
             width_1 = 5.8, #optional, adjust the width of final plot
             folder = "compare_gene_three_lists")
#> [1] "Starting merging gene lists..."
#> [1] "Clustering common gene among three gene lists..."
#> [1] "2 genes with higher frequency in all three gene lists, they are: "
#>      name2_1 Frequency_1 Frequency_2 Frequency_3 Common_Gene
#> 1329   CERS6          21          24          23 Common_High
#> 3361   STK39          21          17          19 Common_High
#> [1] "Brief summary:"
#> # A tibble: 3 x 2
#> # Groups:   Common_Gene [3]
#>   Common_Gene  Number_Gene
#>   <chr>              <int>
#> 1 Common_High            2
#> 2 Common_Low          3788
#> 3 High_and_Low         115
#> [1] "Gene comparison list, brife summary and comparison plot was saved in working directory."
```

Figure.15 Comparison of three gene lists

If we add a fourth gene frequency lists, by adding the `gene_freq_4` argument, the comparison figure produced will be a heat-map rather than a scatterplot.

```
compare_gene(gene_freq_1 = "call_gene_Part/gene_freq_cnv.txt", #standard file was generated by 'call_gene' function in Section 5.2
             gene_freq_2 = "call_gene_Penn_UCSC/gene_freq_cnv.txt", #standard file was generated by 'call_gene' function in Section 5.2
             gene_freq_3 = "call_gene_Part/gene_freq_cnv.txt", #standard file was generated by 'call_gene' function in Section 5.2
             gene_freq_4 = "call_gene_standard_ARS_UCSC/gene_freq_cnv.txt", #standard file was generated by 'call_gene' function in Section 5.2
             title_1 = "Part", #optional, adjust the title of input file 1 present in final plot 
             title_2 = "ENS", #optional, adjust the title of input file 2 present in final plot 
             title_3 = "UCSC", #optional, adjust the title of input file 3 present in final plot 
             title_4 = "PUCSC", #optional, adjust the title of input file 4 present in final plot 
             col_1 = "red", #optional, set the color of High frequency genes
             col_2 = "yellow", #optional, set the color of Low frequency genes
             common_gene_threshold = 20,
             height_1 = 9, #optional, adjust the height of final plot
             width_1 = 11, #optional, adjust the width of final plot
             folder = "compare_gene_four_lists")
#> [1] "Starting merging gene lists..."
#> [1] "Clustering common gene among four gene lists..."
#> [1] "1 genes with higher frequency in all three gene lists, they are: "
#>     name2_1 Frequency_1 Frequency_2 Frequency_3 Frequency_4 Common_check
#> 329   CERS6          21          24          21          23  Common_high
#> [1] "Brief summary:"
#> # A tibble: 3 x 2
#> # Groups:   Common_check [3]
#>   Common_check Number_Gene
#>   <chr>              <int>
#> 1 Common_high            1
#> 2 Common_low          2401
#> 3 High_and_Low          34
#> [1] "Gene comparison list, brife summary and comparison plot was saved in working directory."
```

Figure.16 Heatmap comparing gene lists across four data sets

#### Brief summary

The ten functions we’ve run so far only require users to provide the CNV results, which can already generate plenty of summary results that include sufficient information to point us in the right direction to dig deeper. If we want to explore a particular region in more depth, there are extra functions in the `HandyCNV` package that can combine CNV data with other data sources. We will introduce some of these functions in the following section.

#### 11. How can we create the map file used in `HandyCNV`, to allow comparing CNVs between different reference genomes?

We can use the `convert_map` function to prepare a map file; generally, we will need one for each reference file. In our examples, we have used the two bovine reference genomes ARS-UCD 1.2 and UMD 3.1. For `HandyCNV`, map files hould have the format shown in the table below, with the four columns `Chromosome`, `SNP`, `MorganPos`， and `PhysicalPos`. If the centiMorgan position (`MorganPos`) is unknown, we can assign the missing value as `0`, or provide the phyisical position in megabases. This format is the same as the standard `.map` format used in Plink and other software, and can be exported from GenomeStudio. The column names are not required.

```
tabl <- "
-----------------------------------------
Chr   SNP           MorganPos PhysicalPos
---- ------------- ---------- -----------               
  5   14636         56.3275   56327514 
  5   16084         56.329    56328962  
  5   19597         56.3325   56332475  
  1   1_115292065   114.371   114370731  
  1   1_115292107   114.371   114370773 
-----------------------------------------"
```

Here, we have the map files for both bovine references included in the package, at the following paths:

```
ars_map <- system.file("extdata", "Demo_data/map/ARS1.2_GGPHDV3.map", package = "HandyCNV")
umd_map <- system.file("extdata", "Demo_data/map/UMD3.1_Bovine.map", package = "HandyCNV")
```

The function will not only generate map files, for use in `compare_cnv` and `compare_cnvr`, it will also generate map file used in Plink and PennCNV for both inputs. The conversion process also compares the two input maps and produces summary tables and figures.

```
convert_map(default_map = umd_map, #when running your own data, set this argument to 'default_map = "localpath/file"'
            target_map = ars_map, #when running your own data, set this argument to 'target_map = "localpath/file"'
            chr_set = 30,
            defMap_title = "UMD 3.1", #set the label of axis
            tarMap_title = "ARS-UCD 1.2", #set the label of axis
            col_1 = "green4", #set colour for the type of Match in SNP comparison plot
            col_2 = "red", #set colour for the type of Unmatched in SNP comparison plot
            col_3 = "deeppink2", #set colour for the bar of Target_Map in SNP density plot
            col_4 = "deeppink2", #set colour for line of Target_map in density plot
            col_5 = "turquoise3", #set colour for point of Default_Map in SNP density plot
            col_6 = "turquoise3", #set colour for line of Default_map in SNP density plot
            species = "Bovine", 
            height_c = 16,
            width_c = 22,
            height_d = 14,
            width_d = 25)
#> [1] "Starting comparing the differences between the two versions"
#> [1] "Checking if negtive position exsits in new converted map...."
#> 
#>  FALSE   TRUE 
#> 139512    465 
#> [1] "There are 465 SNPs with negtive postion in new assembly."
#> [1] "The matching results of SNP chromosome ID in the two versions are as follows:"
#> 
#>    Match Mismatch 
#>   122764    17219 
#> [1] "The details of comparison as follows: "
#>                           Type.Var1 Type.Freq
#> 1           Both unknown Chromosome       344
#> 2              Different Chromosome       859
#> 3                   Same Chromosome    122420
#> 4            Unfound in Default map      2038
#> 5             Unfound in Target map         6
#> 6 Unknown Chromosome in Default map     13771
#> 7  Unknown Chromosome in Target map       545
#> [1] "Ploting SNP comparison results....."
#> [1] "Comparison of SNP by Chromosome plot was saved in working directory."
#> [1] "Preparing the map files for PennCNV and Plink...."
#> [1] "Writing converted map for PennCNV and Plink formats..."
#> [1] "Target map was converted to PennCNV map file..."
#> [1] "Converted map for PennCNV format were saved in working directory."
#> [1] "Converted map for Plink format were saved in working directory."
#> [1] "Plotting the Difference of SNPs between two versions of map..."
#> [1] "SNP difference plot was saved in working directory."
```

Figure.17 Comparison of the ARS 1.2 and UMD 3.1 reference genomes

Figure.18 Comparison of SNP densities on the ARS 1.2 and UMD 3.1 maps

#### 12. How can we integrate the CNVs and annotated genes with additional information, such as Log R ratio, B Allele Frequency, call rate, heterozygosity, missing value rate and Linkage Disequilibrium, and plot it as one figure?

The `plot_cnvr_panorama` function was built to achieve this task. But we need provide more information to this function: a. CNVR results(generated by ‘call\_cnvr’ in section 4) b. Annotated CNV file (generated by ‘call\_gene’ function) c. Signal Intensity file (In general it export from GenomeStudio which used in CNV detection process) d. Map file (Four columns used in Plink or generated by ‘convert\_map’ function) e. Genotype files (generated by Plink with ‘–make-bed’ flag which include BED, BIM, FAM three files)

```
plot_cnvr_panorama(cnvr = "call_cnvr_Penn/cnvr.txt", #standard file was generated by 'call_cnvr' function in section 4
                   cnv_annotation = "call_gene_Penn_UCSC/cnv_annotation.txt", #standard file was generated by 'call_gene' function in Section 5.2
                   intensity = "Demo_data/demo_intensity.txt", 
                   map = "Demo_data/demo.map", 
                   prefix_bed = "Demo_data/demo_HandyCNV", 
                   ld_heat = TRUE,
                   sample_size = 382, 
                   common_cnv_threshold = 0.18, 
                   col_0 = "green", #optional, customize color for o copy CNV 
                   col_1 = "hotpink", #optional, customize color for 1 copy CNV 
                   col_2 = "NA", #optional, customize color for 2 copy CNV
                   col_3 = "turquoise", #optional, customize color for 3 copy CNV 
                   col_4 = "aquamarine", #optional, customize color for 4 copy CNV 
                   col_gene = "red", #optional, customize color of gene 
                   width_1 = 16,
                   height_1 = 30,
                   folder = "cnvr_panorama_heat_ld")
#> [1] "4 CNVRs passed the common frequency threshold."
#> Reading Demo_data/demo_HandyCNV.fam 
#> Reading Demo_data/demo_HandyCNV.bim 
#> Reading Demo_data/demo_HandyCNV.bed 
#> ped stats and snps stats have been set. 
#> 'p' has been set. 
#> 'mu' and 'sigma' have been set.
#> [1] "Starting to make plot..."
#> [1] "Plot 1 (7_310937_435584_Mixed.png) was stored in the cnvr_panorama_heat_ld/ directory."
#> [1] "Plot 2 (15_78804694_79611716_Mixed.png) was stored in the cnvr_panorama_heat_ld/ directory."
#> [1] "Plot 3 (12_69804346_70777223_Mixed.png) was stored in the cnvr_panorama_heat_ld/ directory."
#> [1] "Plot 4 (12_56513728_64796304_Mixed.png) was stored in the cnvr_panorama_heat_ld/ directory."
```

Figure.19 Example of CNVR Panorama with LD\_Heatmap

For LD, we can replace the heatmap with the classic inverted triangle diagram by setting the argument `ld_heat = FALSE`, which will use the `gaston` package to plot LD. Because this requires converting the figure to `ggplot` format (via the `ggplotify` package), this process can be time consuming, and sometimes does not work well for large genomic regions. Therefore, we suggest that you use `ld_heat = TRUE` initially, before narrowing in on a particular region of interest to display with the classic LD plot.

Figure.20 Example of CNVR Panorama with Classical\_LD

#### 13. How can we make a plot to show the source of the CNVs?

CNVs are heritable: with the pedigree information, particularly for the sires in a livestock context, we can produce plots of CNV sources. The `plot_cnvr_source` function can produce plots by sire, herd, and sire source. Sire source can be any freeform text, for example the country of origin. The expected input format is comma-separated (with a header), containing the following columns (in any order):

- `Chip_ID` (matched to the `Sample_ID` column in the CNVR data)
- `Herd`
- `Sire_ID`
- `Sire_Source`

```
plot_cnvr_source(cnvr = "call_cnvr_Penn/cnvr.txt", #standard file was generated by 'call_cnvr' function in section 4
                clean_cnv = "cnv_clean/penncnv_clean.cnv", #standard file was generated by 'cnv_clean' function in section 1
                pedigree = system.file("extdata", "Demo_data/Pedigree/Pedigree.csv", package = "HandyCNV"), 
                Frequent_threshold = 80,
                height_1 = 20, 
                width_1 = 15)
```

Figure.21 Plot Source of CNVR

#### 14. How can we plot just the genes in a specific region, to save as a seperate figure?

The `plot_gene` function can do this for a specific reference gene set and genomic region.

```
plot_gene(refgene = "refgene/cow_ARS_UCSC.txt", 
          chr_id = 7, 
          start = 11, 
          end = 12, 
          height_1 = 3, 
          width_1 = 10, 
          show_name = c(11.0, 11.35, 11.75, 12.00), 
          gene_font_size = 2.5)
```

Figure.22 Gene plot

#### 15. How can we find regions with high frequencies of runs of homozygosity?

An ROH (Run of homozygosity) is a region of an individual’s genome where almost all variants have homozygous genotypes. These regions generally indicate the presence of inbreeding in that individual’s ancestry, with more recent inbreeding leading to longer ROHs.

The function `roh_window` will produce two summary tables, as well as plotting the high frequency regions by chromosome. The first table summarises the total numbers and lengths of ROHs for each individual; the second reports summary statistics for aggregated windows of ROHs across all individuals.

```
roh_window(roh = "cnv_clean/cnvpart_roh.cnv", 
           window_size = 2,
           length_autosomal = 2489.386, 
           threshold = 0.15, 
           folder = "roh_window", 
           col_higher = "red", 
           col_lower = "black", 
           height_1 = 15, 
           width_1 = 11, 
           legend_x = 0.92, 
           legend_y = 0.06,
           ncol_1 = 5)
#> [1] "Individual situations are shown below:"
#> # A tibble: 397 x 7
#>    Sample_ID               n total_length min_length max_length means  F_roh
#>    <chr>               <int>        <dbl>      <dbl>      <dbl> <dbl>  <dbl>
#>  1 201094560008_R01C01    56        285.        1.19      21.8   5.08 0.114 
#>  2 201094560008_R01C02    37        176.        1.24      17.4   4.77 0.0708
#>  3 201094560008_R02C01    59        258.        1.11      14.4   4.37 0.104 
#>  4 201094560008_R02C02    35        170.        1.06      15.5   4.86 0.0683
#>  5 201094560008_R03C01    71        324.        1.11      22.2   4.56 0.130 
#>  6 201094560008_R03C02    61        279.        1.03      16.0   4.58 0.112 
#>  7 201094560008_R04C01    91        573.        1.06      26.1   6.29 0.230 
#>  8 201094560008_R04C02    54        376.        1.09      32.1   6.96 0.151 
#>  9 201094560008_R05C01    19         51.1       1.36       9.67  2.69 0.0205
#> 10 201094560008_R05C02    61        336.        1.19      30.7   5.50 0.135 
#> # ... with 387 more rows
#> [1] "The top 10 ROHs by frequency are as follows:"
#>    window_id number_roh Chr Start_win End_win prop_roh
#> 1    Win_355        220   6   3.8e+07 4.0e+07    0.554
#> 2    Win_649        170  11   6.6e+07 6.8e+07    0.428
#> 3    Win_354        150   6   3.6e+07 3.8e+07    0.378
#> 4    Win_382        143   6   9.2e+07 9.4e+07    0.360
#> 5    Win_381        142   6   9.0e+07 9.2e+07    0.358
#> 6    Win_366        135   6   6.0e+07 6.2e+07    0.340
#> 7    Win_380        133   6   8.8e+07 9.0e+07    0.335
#> 8    Win_650        132  11   6.8e+07 7.0e+07    0.332
#> 9    Win_648        131  11   6.4e+07 6.6e+07    0.330
#> 10   Win_859        123  16   2.8e+07 3.0e+07    0.310
#> [1] "Plotting frequncy of ROH by windows..."
#> [1] "Task done."
```

Figure.23 ROH Frequency Distribution

#### 16. How do we visualise ROHs?

Visualising ROH is similar to visualising CNVs. The exception is that, because length is particularly informative for ROHs (about the age of inbreeding), the ROHs are coloured by their lengths. The `roh_visual` function can plot ROHs, either across the whole genome, or chromosome by chromosome. It also supports restricting plotting to regions of interest, and allows adding a panel of gene locations.

##### 16.1 Plot ROH by Chromsome

To plot a single chromosome, use the `chr_id` argument.

```
roh_visual(clean_roh = "cnv_clean/cnvpart_roh.cnv", 
           chr_id = 6, 
           width_1 = 14, 
           height_1 = 9, 
           folder = "roh_visual", 
           col_short = "red", 
           col_mid = "blue", 
           col_long = "black")
#> [1] "Preparing plot data..."
#> [1] "Task done, plot was stored in working directory."
```

Figure.24 ROH on Chromosome 6

##### 16.2 Plot ROH by single region

To plot a single region of interest, add the `start_position` and `end_position` arguments (units in megabases).

```
roh_visual(clean_roh = "cnv_clean/cnvpart_roh.cnv", 
           chr_id = 6, 
           start_position = 36.5, 
           end_position = 40.6, 
           width_1 = 16)
#> [1] "Preparing plot data..."
#> [1] "Task done, plot was stored in working directory."
```

Figure.25 ROH on Chr6:35.0-42.5 Mb

##### 16.3 How can we present annotated genes above the ROH plot?

To add a panel of gene locations, add the `refgene` argument, pointing to a file containing genes annotated on the appropriate reference genome.

```
roh_visual(clean_roh = "cnv_clean/cnvpart_roh.cnv", 
           chr_id = 6, 
           start_position = 36.5, 
           end_position = 40.61, 
           refgene = "refgene/cow_UMD_UCSC.txt")
#> Warning: Removed 2 rows containing missing values (geom_vline).
```

Figure.26 Annotation of ROH on Chr6:36.5-40.5

If the ROH region has too many genes, plotting all the gene names will make the figure look too busy. We can choose the most important region(s) to plot using the `show_names` argument. In this example, we set two regions with the following argument: `show_name = c(37.28, 37.49, 38.25, 39.52)`. This will show the gene names in two regions: (37.28 - 37.49 Mb) and (38.25 - 39.52 Mb).

```
roh_visual(clean_roh = "cnv_clean/cnvpart_roh.cnv", 
           chr_id = 6, 
           start_position = 36.5, 
           end_position = 40.5, 
           refgene = "refgene/cow_UMD_UCSC.txt", 
           show_name = c(37.28, 37.49, 38.25, 39.52), col_short = "black", col_mid = "green", col_long = "red")
```

Figure.27 Customised gene annotations on Chr6:36.5-40.5 Mb

##### 16.4 How do we plot the full length of ROH for all samples that had a ROH in a given region?

Above, We have plotted ROH in an interested region, but that method can only plot short ROHs which located in the given region, it will remove some longer ROHs which are partially overlapped to the given interval, that method can made a nice picture but will lost some samples, it still useful because we already know the sample size in that region. But if we want to present the whole length of ROH overlapped to our interested region (Such as a region of target gene), we can assign a vector to ‘target\_region’ argument in ‘roh\_visual’ function.

```
roh_visual(clean_roh = "cnv_clean/cnvpart_roh.cnv", 
           target_region = c(6,38.25,39.52), #within the vector, the first integer indicate Chromosome. the rest are the start and end point, unit in megabases
           col_short = "red", #optional
           col_mid = "gold", #optional
           col_long = "blue", #optional
           height_1 = 12, #optional
           width_1 = 14)
#> [1] "Preparing plot data..."
#> [1] "Task done, plot was stored in working directory."
```

Figure.28 ROHs with full length on target region Chr6:38.25-39.52 Mb

##### 16.5 How do we plot ROHs on an interested sample?

When we have some interested individuals, we can plot all the ROHs for these samples separately by setting the ‘individual\_id’ arguments as following:

```
roh_visual(clean_roh = "cnv_clean/cnvpart_roh.cnv", 
           individual_id = "201094560008_R04C01", 
           height_1 = 14, #optional 
           width_1 = 22,  #optional
           col_short = "green", #optional
           col_mid = "yellow", #optional
           col_long = "red")
#> [1] "Preparing plot data..."
#> [1] "Task done, plot was stored in working directory."
```

Figure 29. ROHs on interested Sample

#### 17. How do we get haplotype for ROH region?

ROH results will return us some long regions, but we may interested on some specific genes in the ROH region that experienced strong selection. Get the haplotypes of these genes in our experimental population may help us to find out some interesting patterns. we known that the ROH region generated by SNPs may be discontinuous due to insufficient marker density, beside, when infer ROH we considered the influence of genotyping quality by setting the minimal window size, accepting some missing value or maximum heterozygosity SNPs in the window, these conditions will exclude the short ROH regions in the final results. For instance, when we get the haplotype for a short region (such as a Gene), the frequency of homozygosity haplotypes may differing to the original ROH region it located.

The ‘get\_haplotype’ function can help us to get the haplotype for all samples by given region. But now the application is limited, it only accept the phased genotype by Beagle (Version 5.1) with physical position, we may expend the compatibility later if necessary. The standard format of phased genotype and demo data used at here can be download from our github repository (https://github.com/JH-Zhou/HandyCNV/tree/master/vignettes/OARS\_chr6). Now, let’s go ahead and make haplotypes.

Extracting the haplotype will take three steps and combine three functions. The first step is to prepare the standard format for ‘HandyCNV’. We can use ‘prepare\_geno’ function to prepare the phased genotype. After running, it will return three list files contain the map, genotype and the columns index file.

```
genotype <- prep_phased(phased_geno = "OARS_chr6.vcf") #'OARS_chr6.vcf' is the demo data, need to download and save it in your working directory 
#> [1] "Reading input file..."
#> [1] "Spliting Sample columns..."
#> [1] "Tansposing genotype..."
```

Second step, we need to assign the interested region. Here we take the *NCAPG* gene as example, it located on BTA6:37,332,024-37,378,129 bp. The idea in this step is to find out the closest SNPs in this SNP chip.

```
region <- closer_snp(phased_input = genotype$map, #the map file in the list generated in 'pre_phased' function above
                     chr = 6, start = 37.332024, end = 37.378129) #physical position unit in 'megabases'
#>        min      max
#> 1 37340937 37377606
```

Third step to get haplotype. It will return two lists contains the reocding haplotype and the index of recode type and the original haploid.

```
haplotype <- get_haplotype(geno = genotype, pos = region)
#> [1] 2481
#> [1] 2490
#> [1] "Recoding Haploid..."
#> [1] "Generating Haplotype"
```

Now, we can count the frequency of each haplotype in our population and to check what the haploids are in these haplotypes.

```
data.frame(table(haplotype$hap_type$Recode_Type, dnn = "Haplotype"))
#>    Haplotype Freq
#> 1       1011    1
#> 2         11  353
#> 3         12   26
#> 4         14    5
#> 5         19    1
#> 6         22    1
#> 7         33    6
#> 8         55    1
#> 9         66    1
#> 10        77    1
#> 11        88    1
haplotype$hap_index
#> # A tibble: 11 x 3
#>    Haploid     Freq Re_code
#>    <chr>      <int>   <dbl>
#>  1 0000000000   738       1
#>  2 1111111111    28       2
#>  3 000000N000    12       3
#>  4 1111101100     5       4
#>  5 00000N0000     2       5
#>  6 0N00000000     2       6
#>  7 0N0000N000     2       7
#>  8 N000000000     2       8
#>  9 0000001000     1       9
#> 10 0N0000000N     1      10
#> 11 0N0000100N     1      11
```

#### 18. How do we convert coordinates for CNV, CNVR, ROH, or any other intervals?

The function `convert_coord` can used to convert coordinates for lists of intervals. Currently, the function is limited to inputs generated by the `convert_map` function, and can only convert the coordinates for intervals on the same type of SNP Chip. Converting coordinates may change the total length of the intervals, as of the positions and orders of the SNPs on the chromosome will potentially differ between various versions of reference genome; therefore, we should pay attention to the quality of the conversion, and use the results with caution. The function produces a summary table that summarises how many intervals were converted successfully, and reports on the differences in length between the converted and original intervals.

The `proxy_range` argument supports adjusting the genomic range to search and assign “proxy” locations for markers which are missing in the target map, in order to try to best reduce the loss of intervals after conversion. The principal is try to reduce the length of the intervals that were assigned to the proxy locations. For missing markers at the start positions, the search will begin at the missing position and proceed downsteam, selecting the closest suitable marker as the proxy position. For the end position, the search will begin from the missing end interval and proceed upsteam, again assigning the closest suitable marker as the proxy position. If no suitable markers can be found within `proxy_range` basepairs, the interval will be classed at “Missing”.

The optional argument `len_diff_threshold` is a quality control measure on the newly converted intervals that imposes a maximum difference of length between converted and original intervals. Only intervals with remapped positions within `len_diff_threshold` basepairs of the original position will pass the threshold.

Here, we convert the CNVPartition CNVR results from UMD3.1 coordinates to ARS1.2. The outputs are saved into the `convert_coord_penn_cnv` folder, and summary statistics about the conversion are displayed below.

```
convert_coord(input_def = "cnv_clean/cleancnv.cnv", 
              def_tar_map = "convert_map/def_tar_map.map",
              proxy_range = 100000,
              len_diff_threshold = 100000,
              folder = "convert_coord_penn_cnv")
#> [1] "'def_tar_map' file passed input check..."
#> [1] "The quality check of converion on Start Position as below: FALSE means not match between two version."
#> 
#>    Match Mismatch 
#>     4396      180 
#> [1] "The quality check of converion on End Position as below: FALSE means not match between two version."
#> 
#>    Match Mismatch 
#>     4322      254 
#> [1] "4104 Intervals passed quality check."
#> 
#> Fail Pass 
#>  472 4104 
#>                                  Item    Length Number
#> 1                      Total Original 974963676   4576
#> 2                   Succeed Original  891042504   4104
#> 3                  Succeed Convertion 897762659      -
#> 4                   Failed convertion  77201017    472
#> 5 Difference Converted minus Original  -6720155      -
```

We can also convert coordinates from the target map to the default map. For example, let’s convert the PennCNV ARS results to UMD 3.1. After converting the coordinates of CNVs or CNVRs, we could use the results which passed the quality check to make comparisons with the `compare_cnv` and `compare_cnvr` functions as section 8 and 9 described.

```
convert_coord(input_def = "call_cnvr_standard/cnvr.txt", 
              proxy_range = 100000,
              len_diff_threshold = 100000,
              def_tar_map = "convert_map/def_tar_map.map", 
              folder = "convert_coord_penn_cnvR")
#> [1] "'def_tar_map' file passed input check..."
#> [1] "The quality check of converion on Start Position as below: FALSE means not match between two version."
#> 
#>    Match Mismatch 
#>      871       10 
#> [1] "The quality check of converion on End Position as below: FALSE means not match between two version."
#> 
#>    Match Mismatch 
#>      870       11 
#> [1] "835 Intervals passed quality check."
#> 
#> Fail Pass 
#>   46  835 
#>                                  Item    Length Number
#> 1                      Total Original 286942723    881
#> 2                   Succeed Original  270103909    835
#> 3                  Succeed Convertion 271977583      -
#> 4                   Failed convertion  14965140     46
#> 5 Difference Converted minus Original  -1873674      -
```
